# Precision-cut lung slices in air-liquid interface (PCLS-ALI): A novel *ex-vivo* model for the study of Pulmonary Aspergillosis

**DOI:** 10.1101/2024.11.15.615211

**Authors:** LE Gonzales-Huerta, TJ Williams, R Aljohani, BD Robertson, CA Evans, DPH Armstrong-James

## Abstract

Pulmonary Aspergillosis is a respiratory infection with a high mortality rate, which affects patients with immunosuppression or structural lung defects. Antifungal treatment options are few and many have narrow therapeutic margins and potentially serious side effects. In recent years, there are growing numbers of reports of antifungal resistance. Thus, there is an urgent need for effective models to study fungal pathogenesis and test antifungal therapies in the respiratory system. Here, we present a novel *ex-vivo* model using precision-cut lung slices in an air-liquid interface platform to evaluate lung tissue responses to fungal infection and antifungal treatment. Readouts assessed were lactate dehydrogenase for tissue damage, release of inflammatory cytokines (TNF-α, IL-1β, CXCL1), and histology for confirmation of hyphal invasion. Overall, the PCLS-ALI model is a promising approach for understanding lung tissue responses to fungal infections, which fulfils the reduction and refinement components of the 3Rs guiding principles for ethical use of experimental animals.

## INTRODUCTION

Pulmonary aspergillosis (PA) is primarily caused by *Aspergillus fumigatus* (*Af*) which is found ubiquitously in the air and easily reaches the lower respiratory tract. Those with high susceptibility to PA can be broadly categorized within two groups, first those with dysfunctional immune systems including immunodeficiency, immunosuppression, or hypersensitivity and, secondly, those with structural lung disease such as bronchiectasis or pulmonary cavitations (1). PA is strongly associated with neutropenia, which makes patients with cancer more susceptible to this infection (2). Most antifungal drugs routinely used clinically, including Amphotericin B (AmB) (3,4) and the triazoles, have toxic side effects. These issues, together with increasing reports of antifungal resistant strains of *Af*, both clinical and environmental, highlight a need for models or the investigation of antifungal drugs in a respiratory setting (5). To experimentally replicate immunosuppression and generate a PA animal model, pharmacological immunosuppression is typically used (6). However, these models are time consuming, use large numbers of animals, can expose animals to repeated procedures over long periods of time, and lack the diverse remodelling of lung tissue seen in human pathology (7–10). Precision-cut lung slices (PCLS) have been used since the late 1980s, when Placke and Fisher optimised the infusion of low-melting point agarose into the lungs, allowing their slicing without damaging the structural organization of the tissue. PCLS have been shown to be suitable for several downstream applications investigating immune responses including flow cytometry, microscopy, quantification of cytokine and chemokine release, and RNA isolation (11). Recently, this *ex-vivo* model has enabled the study of viral and bacterial infections and the efficacy of new drugs on a larger scale compared to *in-vivo* experiments (12) (13) (14). Nonetheless, to our knowledge, respiratory fungal pathogens have not been tested using this model.

Air-liquid interface (ALI) models have been widely utilised for investigating processes that occur in the lung between epithelial layers and immune cells. The principle is to provide two environments in one cell culture model, with a layer submerged in medium and another layer exposed to air to simulate some of the histological conditions involved in gas exchange (15). Additionally, it is suggested that ALI models are superior to traditional submerged cultures of cells since they better replicate the conditions involved in surfactant production and cell migration (16). ALI models have proven valuable to test toxicity of substances and drugs, as well as the delivery of aerosolized substances (17). This model has been successfully tested using cell lines and primary cells, mainly focused on a respiratory epithelium model; nonetheless, co-culture systems that include macrophages, dendritic cells and endothelial cells have also been employed (18).

In the context of *Af* infection, the ALI model offers an additional benefit by modelling the interface where fungal invasion occurs. Most infection models, either *in-vitro* or *ex-vivo*, submerge cells or tissue in culture medium, requiring the submersion of the pathogen tested. This is an additional limitation when working with *Af*, since it adapts efficiently to the nutrients available and its behaviour can vary significantly accordingly (19).

Here we describe the design and performance of a system that combines PCLS and ALI models to create a novel, physiologically relevant, *ex-vivo* model for the study of lung respiratory fungal infections, which simultaneously complies with the principles of the 3Rs for humane animal research, reducing the number of experimental animals in comparison with conventional approaches.

## METHODS

### Ethics and animal use

All mouse work was carried out following the Animal [Scientific Procedures] Act 1986. Procedures were reviewed by the Imperial College London Animal Ethics Committee and performed in accordance with the project licence PPL PF9534064 and personal licence PPLI90527671. Healthy C57BL/6J mice between 8 and 12 weeks old were euthanized with intraperitoneal administration of 0.2 mL of pentobarbital. Secondary confirmation of death was performed by dissection of the femoral blood vessels. Cadavers were immediately used for experimentation.

### Lung extraction

Mice were sprayed liberally with 70% ethanol to prevent contamination. A mid-line dissection from the middle of the abdomen to the neck was performed, followed by thoracotomy to expose heart, lungs, and trachea. A small cut in the ventral side of the trachea, proximal to the cricoid cartilage, was performed. 1 mL of warm (37-40°C) 2% agarose low-melting point (Promega® Cat. V2111) was infused through the trachea into the lungs using a polythene tube with a 1.5 mm diameter (Portex Cat. 800/100/280) connected to a 1 mL syringe. Once lungs were fully inflated, the mouse was covered with aluminium foil and ice put on top for 10 min. Once lungs were full and solidified, both lungs and heart were dissected out of the thoracic cavity *en bloc* and immersed in RPMI-1640 (Gibco® Cat. 72400-047) + 5 μg/mL of Gentamicin (Gibco® Cat. 15750-060) at 4°C.

### Precision cut-lung slices (PCLS)

The tissue bath of a 5100 mz vibrotome (Campden Instruments®) was filled with cold HBSS (Gibco® Cat. 14025-092). Lung lobes were dissected and adhered to the specimen plate with superglue. For slicing, the vibrotome settings were as follow: frequency 80 Hz, amplitude 1.50 mm, section 500 μm, blade speed 0.50 – 0.80 mm/s. Slices with shortest length over 5 mm were considered suitable for experimentation and transferred to petri-dishes with RPMI-1640 (Gibco® Cat. 72400-047) + 10% heat-inactivated foetal bovine serum (FBS) (Gibco® Cat. 10500-064) + 5 μg/mL of Gentamicin (Gibco® Cat. 15750-060). Slices were incubated overnight at 37°C + 5% CO_2_ to allow agarose to dissolve. The following day, lung discs were obtained from slices using a 5 mm diameter biopsy punch instrument. Lung discs were transferred to a 96-well tissue culture plate or HTS Transwell® 96-well plate (Corning® Cat. 3384).

### Selection of culture medium

For assessing medium acidification, phenol red absorbance at 415 nm was measured from lung discs incubated in a 96-well tissue culture plate at 37°C + 5% CO_2_ for 48 hours in RPMI-1640 w/phenol red + 10% heat-inactivated FBS + 5 μg/mL of Gentamicin.

To test the effect of the type of medium on inflammatory cytokine release, we measured TNF-α and IL-1β in the supernatant from lung discs cultured with RPMI-1640 + 10% FBS or RPMI-1640 + 1% FBS or OptiMEM® + 1% FBS. For TNF-α, lung discs were stimulated for 4 hours with LPS [25 ng/mL] (Sigma® Cat. L2630). For IL-1β, discs were stimulated for 3 hours with LPS [25 ng/mL] + 45 min with Nigericin sodium salt [20 μM] (Sigma® Cat. N7143-5MG). Supernatants were then collected and stored at -80°C until tested Bio-Techne ELISA kit for TNF-α (R&D Systems Cat. DY410) and IL-1β (R&D Systems Cat. DY401).

### Culture and preparation of *Aspergillus fumigatus*

*Aspergillus fumigatus* of the CEA-10 strain was cultured in Sabouraud dextrose agar (Oxiod® Cat. CM0041B) in T25 non-ventilated flasks for 3-4 days at 37°C. On the day of infection, fungi were harvested by washing the surface of the agar with DPBS (Gibco® Cat. 14190-136) + 0.1% Tween20 and filtering the conidia suspension through sterile Miracloth™. Fungi were centrifuged at 3000 x g for 10 min, the supernatant was discarded, and fungi resuspended in culture medium and enumerated using a haemocytometer.

### Infection of lung tissue in an air-liquid interface (ALI) model

Lung discs were placed on top of the permeable support of HTS Transwell® 96-well plates (Corning® Cat. 3384) containing 200 μL of Opti-MEM® (Gibco® Cat. 11058-021) + 1% FBS, without antibiotics, and immediately infected by adding 10^5^ resting CEA-10 conidia resuspended in 50 μL of Opti-MEM® + 1% FBS on top of the air exposed tissue. The plate was immediately incubated at 37°C + 5% CO_2_ for 48 h.

### Antifungal assay

For antifungal testing, culture medium with Amphotericin B (Sigma® Cat. A2942-100ML) at 0.1 μg/mL, 1 μg/mL, and 2 μg/mL was added into the wells prior to the introduction of lung tissue discs.

### LDH assay

LDH release was quantified using the Cytotox-96® Non-Radioactive Cytotoxicity Assay (Promega® Cat. G1780) according to manufacturer’s instructions. For the standard curve, a titration was performed with L-Lactate Dehydrogenase (Roche Diagnostics GmbH Cat. 10127230001).

### ELISA for inflammatory cytokines

Cytokines were quantified using a sandwich ELISA kit from Bio-Techne for TNF-α (R&D Systems Cat. DY410), IL-1β (R&D Systems Cat. DY401) and CXCL1/KC (R&D Systems Cat. DY453) according to manufacturer’s instructions. For CXCL1, supernatant was diluted 1:100 using 1% BSA in DPBS.

### Histology

Lung discs were removed gently from the permeable support and fixed with 4% paraformaldehyde (PFA) for 2 hours, washed once with PBS, then transferred to a tissue embedding cassette, and submerged in 70% ethanol prior to sectioning. Next, tissue was wax embedded and sectioned transversally to 10 μm and stained with haematoxylin/eosin (H&E). Images were taken using a Zeiss wide-field microscope at 200x magnification.

### Statistical analysis

All statistical analysis were performed on biological replicates. To determine statistical significance between two unmatched groups, Student’s-T test was applied. For statistical significance, in data normally distributed, between more than two groups in one variable, the One-way analysis of variance (ANOVA) was applied. Comparisons between two or more groups and two variables were analysed using Two-way ANOVA. Identification of outlier values was performed using Grubbs’s test. Statistical analysis was perform using GraphPad Prism 9.4.1.

## RESULTS

### Optimisation for the generation of lung discs from precision-cut lung slices (PCLS)

Medium acidification may interfere with the accurate interpretation of results, since pH has been shown to modulate inflammatory cytokine release in mammalian cells (20), as well as fungal growth and virulence (21,22). Phenol Red absorbance at 415 nm is a reliable method for assessing medium pH. As pH decreases, absorbance at 415 nm increases, reaching its peak value at pH < 5.5 (23,24). Mean values, standard deviation (σ) and margins of error were calculated for confidence intervals (CI) of 95%.

**Schematic 1.**
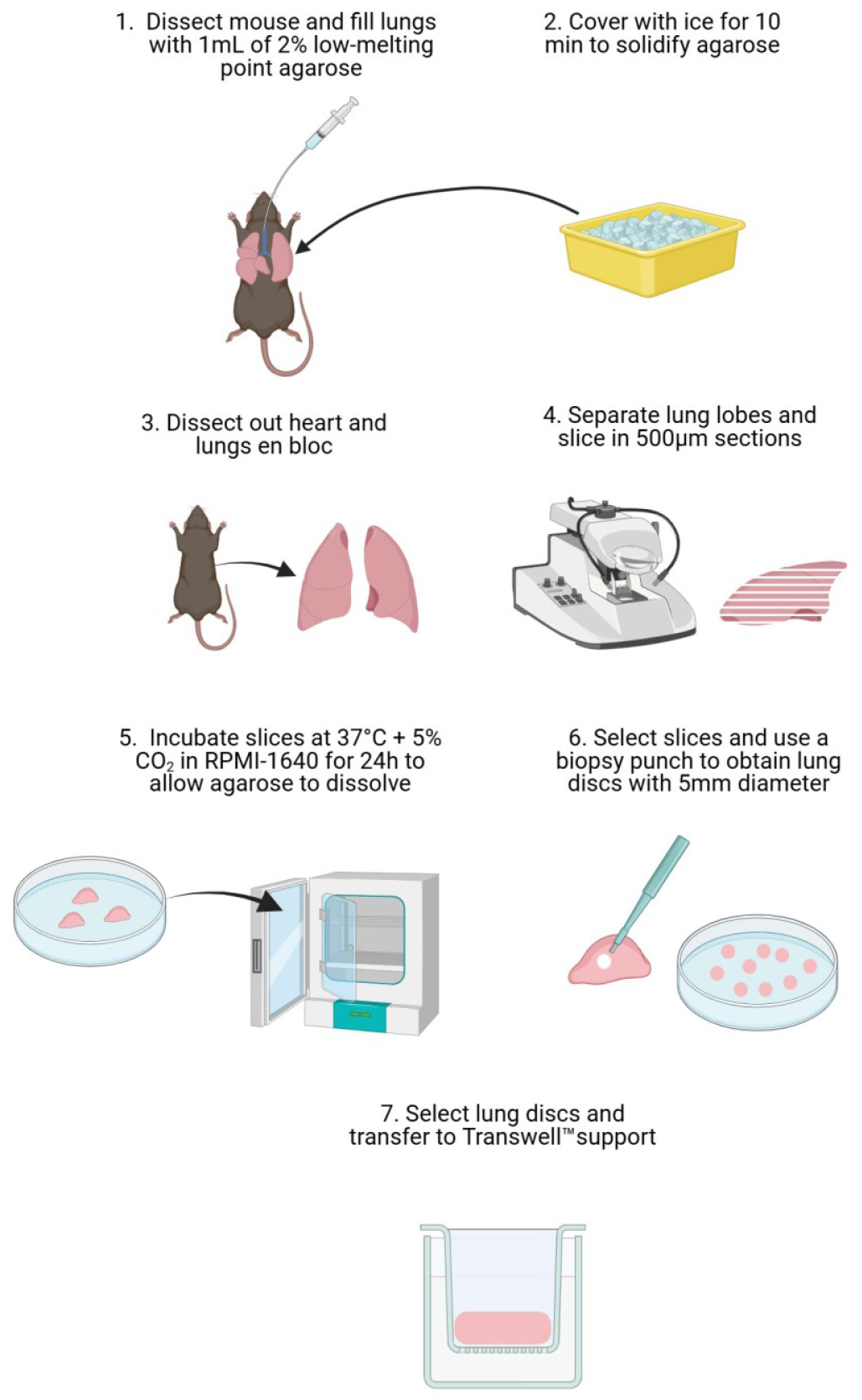
Steps for generating lung discs from PCLS. **1**. C57BL/6J mice were euthanized via intraperitoneal administration of 0.2mL of pentobarbital. Secondary confirmation was performed with dissection of the femoral vessels. Careful dissection of the neck and thoracic structures was performed, and trachea was cannulated for infusion of 1mL of 2% low-melting point agarose. **2**. Lungs were covered with aluminium foil and ice is added over it for 10min to solidify agarose. **3**. Lungs and heart are dissected and extracted carefully en bloc. **4**. Lungs were dissected into individual lobes, which were stuck on the vibrotome’s specimen holder with superglue and sliced in sections of 500μm thickness. **5**. Lung slices were transferred to petri-dishes with culture medium and incubated for 24 hours. **6**. Slices with adequate dimensions were selected and a surgical biopsy punch was used to generate lung discs of 5mm diameter. **7**. Discs with adequate tissue integrity and regular borders were selected for experimental work

**Photographs.**
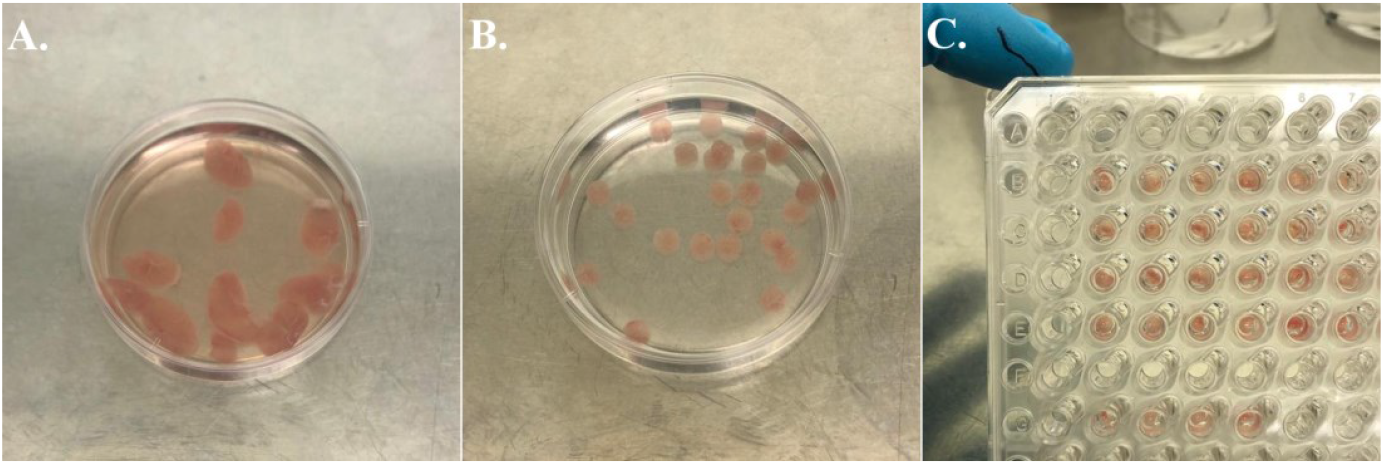
**A**. Petri dish with fresh lung slices in culture medium. A bright pink colour is a sign of healthy tissue. **B**. Lung discs obtained with a biopsy punch were selected for having homogenous, smooth borders and keep bright pink colour after 24 hours of incubation. **C**. Lung discs fit properly on the HTS Transwell® 96-well plate support.

In the absence of tissue, the variability, attributed to incubation and equipment readings, ranged below 5%. For single lung discs, we obtained an absorbance (Abs) mean value of 0.149, with a standard deviation (σ) of 0.023, and margins of error of ± 5.94% for CI = 95%. Increasing the amount of tissue per well, by adding a second lung disc, yielded an Abs mean value of 0.212, a σ=0.052 and margins of error of ± 16.05%. Three lung discs per well reduce the variability, showing a Abs mean value of 0.253, a σ=0.032 and margins of error of ± 8.21%. Statistical analysis showed differences between medium versus 2 lung discs (p=0.006), medium versus 3 lung discs (p=0.001), and 1 lung disc versus 3 lung discs (p=0.012). This showed that the processing of the tissue produced heterogeneous lung discs that avoided significant differences in culture medium acidification (Fig. 1).

**Figure 1.**
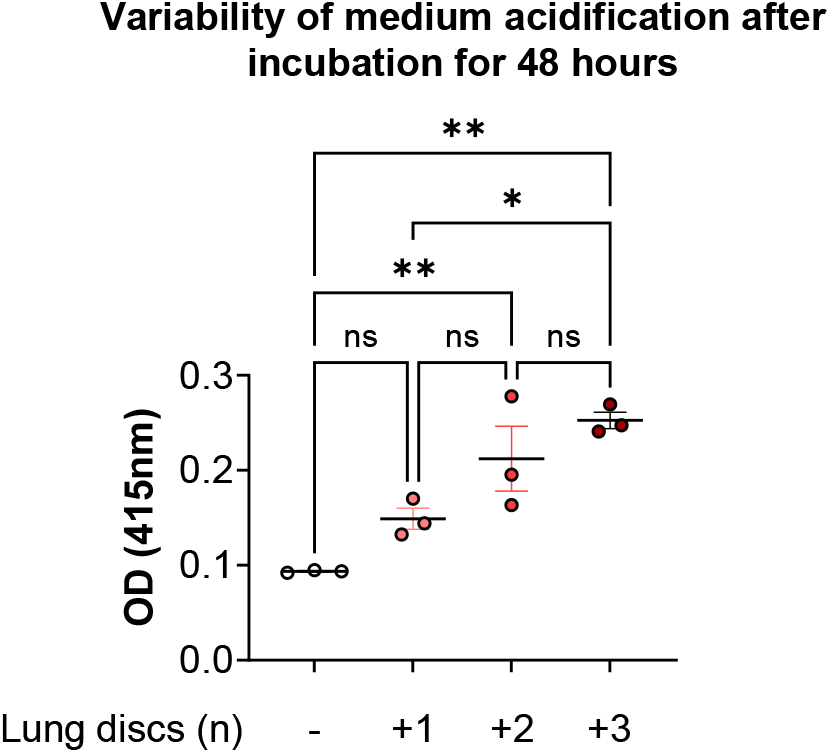
Effect of lung discs heterogeneity in medium acidification. Lung tissue discs were cultured in RPMI-1640 + 10% FBS, 2 mM GlutaMAX®, 25 mM HEPES and readout was taken after 48 hours of incubation at 37°C in 5% CO_2_. Experimental design included technical triplicates. Statistical significance tested with one-way ANOVA on biological replicates (n=3). p value * < 0.05, ** < 0.01, *** < 0.001, **** < 0.0001.

### OptiMEM®+1% FBS benefits TNF-α release upon stimulation with LPS

We investigated the impact culture medium had on cell and tissue viability, comparing OptiMEM®+1% FBS, RPMI-1640+1% FBS, and RPMI-1640+10% FBS. Cell viability was assessed by measuring the LDH in the supernatant after 48 hours. No significant differences in LDH release were detected; however, OptiMEM®+1% FBS showed the lowest standard deviation with σ=14.142 versus σ=20.915 and σ=16.240 for RPMI-1640+1% FBS and RPMI-1640+10% FBS respectively (Fig. 2 B).

**Figure 2.**
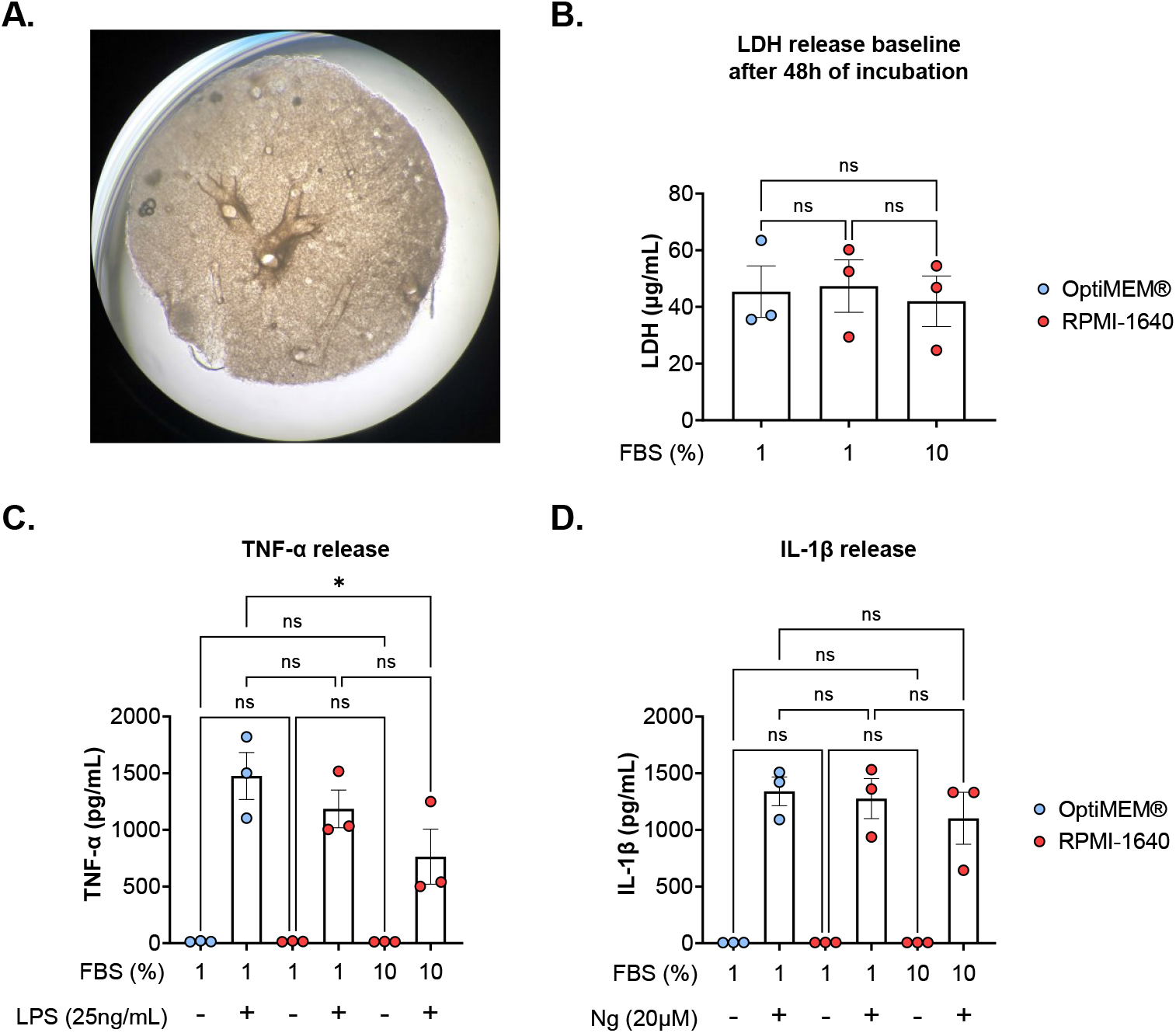
OptiMEM® provides the strongest TNF-α response under stimulation. **A**. Representative photograph of lung disc in culture plate. **B**. Comparison between the effects of OptiMEM® + 1% FBS, RPMI-1640 + 1% FBS and RPMI-1640 + 10% FBS on the LDH baseline from lung discs after 48 hours without stimulation. **C**. Comparison between the effect of 3 different culture medium on the release of TNF-α after 4 hours of LPS [25 ng/mL] stimulation shows stronger response with OptiMEM® + 1% FBS. **D**. Comparison between the effect of 3 different culture medium on the release of IL-1β after stimulation with Nigericin [20 mM]. One-way ANOVA on biological replicates (n=3).

Next, we assessed how the culture medium alters the inflammatory response of the tissue by measuring TNF-α and IL-1β release in response to inflammatory stimuli. At resting state there was no TNF-α or IL-1β release detectable from tissue discs after 48 hours. Upon stimulation with LPS, lung discs in OptiMEM®+1% FBS yielded the highest levels of TNF-α, 1475 pg/mL (210-2733 pg/mL), similar to that of tissue discs cultured in RPMI-1640+1% FBS, 1185 pg/mL (783-1604 pg/mL), but significantly greater than that of lung discs cultured in RPMI-1640+10% FBS, 764 pg/mL (142-1413 pg/mL) (Fig. 2 C).

After stimulation with nigericin, lung discs cultured with OptiMEM®+1% FBS had a mean concentration of 1340 pg/mL IL-1β (1060-1535 pg/mL). For RPMI-1640+1% FBS, IL-1β level was 1277pg/mL (896-1539pg/mL), and for RPMI-1640+10% FBS, 1102 pg/mL (868-1492 pg/mL, with 1 outlier at 26 pg/mL) (Fig. 2 D).

### *Af* infection induces LDH and TNF-α release

To determine the fungal burden required to obtain a measurable tissue response, lung discs were infected with 3 different doses of *Af* (1,000, 10,000, and 100,000 conidia) for 48 hours and the release of LDH and TNF-α was quantified in the supernatant. Mycelial-phase growth of *Af* was evident on top of the lung disc fully covering the air exposed side of the interface (Fig. 3 A, B). Histological H&E staining of tissue discs showed hyphal growth and infiltration into the lung tissue (Fig. 3 D). LDH release from tissue discs was significantly increased when infected with live *Af* conidia. For uninfected lung discs, the mean concentration of LDH was 104 μg/mL (61-168 μg/mL). For infected discs, the mean concentrations of LDH were 135 μg/mL (124-167 μg/mL) for 1,000 conidia, 154 μg/mL (95-188 μg/mL) for 10,000 conidia, and 155 μg/mL (123-173 μg/mL) for 100,000 conidia (Fig. 3 E).

**Figure 3.**
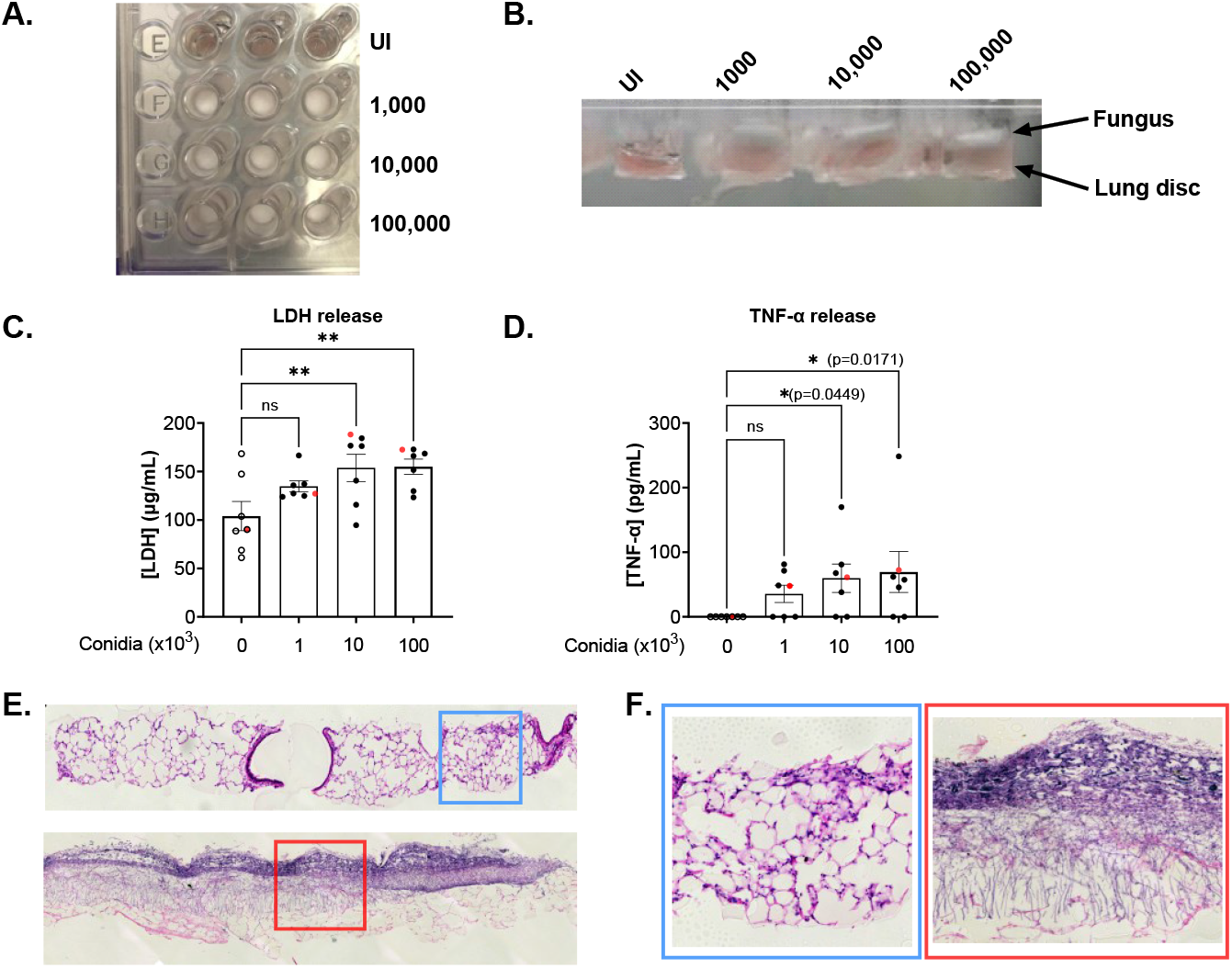
Release of LDH and TNF-α is dependent on fungal burden. **A**. Representative photograph of lung discs on Transwell™ supports after 48 hours post infection with different concentrations of *Af*. **B**. Side view of Transwell™ supports showing air-liquid interface and fungus growing predominantly in the air section as a white, mycelial layer. **C**. LDH released to supernatant 48hpi. **D**. Concentration of TNF-α in supernatant 48 hpi. Data points in red represent values obtained from wells shown in panels A and B. **E**. Representative images of an uninfected (top) and an infected (below) lung disc with 100,000 *Af* conidia for 48 hours (H&E staining, x20). For uninfected conditions, the histological characteristic of the lung is preserved after 48 hours of incubation on ALI. Infected lung discs show fungal layer with infiltration of hyphae. **F**. Zoom image (x10) from panel E. (blue = Uninfected, red = Infected). Statistical significance was tested with one-way ANOVA on biological replicates (n=7). p value * < 0.05, ** < 0.01, *** < 0.001, **** < 0.0001

For TNF-α, all uninfected lung discs showed values below the limit of detection (LOD). Infected lung discs had a mean concentration of TNF-α of 36 pg/mL (0-81 pg/mL) for 1,000 conidia, 60 pg/mL (0-170 pg/mL) for 10,000 conidia, and 69 pg/mL (0-73 pg/mL, outlier at 248 pg/mL) for 100,000 conidia (Fig. 3 F). Statistically significant differences were found for uninfected versus 10,000 conidia (p=0.045) and 100,000 conidia (p=0.017). Despite the absence of noticeable macroscopical difference between the three doses, 100,000 conidia showed the greatest release of LDH with less variability between samples, therefore, this dose was used for subsequent experiments.

### Amphotericin B (AmB) restricts *Af* growth and fungal infiltration into tissue, leading to reduced inflammation in the PCLS-ALI model

We used the optimised ALI model of infection to investigate antifungal drug efficacy. We tested AmB, a powerful antifungal with a narrow therapeutic margin (25), using a dose titration designed to cover a range from subtherapeutic (0.1 μg/mL) to toxic concentrations (2 μg/mL).

At 48 hours post infection (hpi), uninfected lung discs, 2 μg/mL of AmB has no statistically significant effect on cell viability and inflammatory cytokines release (Fig. 4 A B C). In infected lung discs, AmB restricted fungal growth in a dose dependent manner (Fig. 4 D), which correlated with the concentrations of LDH in the supernatant (Fig. 4 E). Furthermore, AmB modulated inflammatory cytokine release. Although, no significant differences for TNF-α were detected, IL-1β showed a strong, dose-depended response. IL-1 β increased from 24 pg/mL in untreated tissue to 60 pg/mL with 2 μg/mL of AmB (Fig. 4 F and G). Additionally, AmB activity against fungal growth correlated with concentrations of CXCL1 in supernatant. Baseline concentration of CXCL1 in the supernatant of lung discs without infection and treatment were considerably high and required serial dilutions to test successfully. However, *Af* infection reduced CXCL1significantly, from 77 ng/mL in uninfected, untreated tissue, to 22 ng/mL in infected, untreated lung discs. AmB treatment was associated to CXCL1 increase in a dose-dependent manner (Fig. 4 H). Histological sections showed that *Af* hyphae infiltrated effectively into the lung discs and that AmB at 1 μg/mL or higher concentrations severely disrupted fungal morphology (Fig. 4 I).

**Figure 4.**
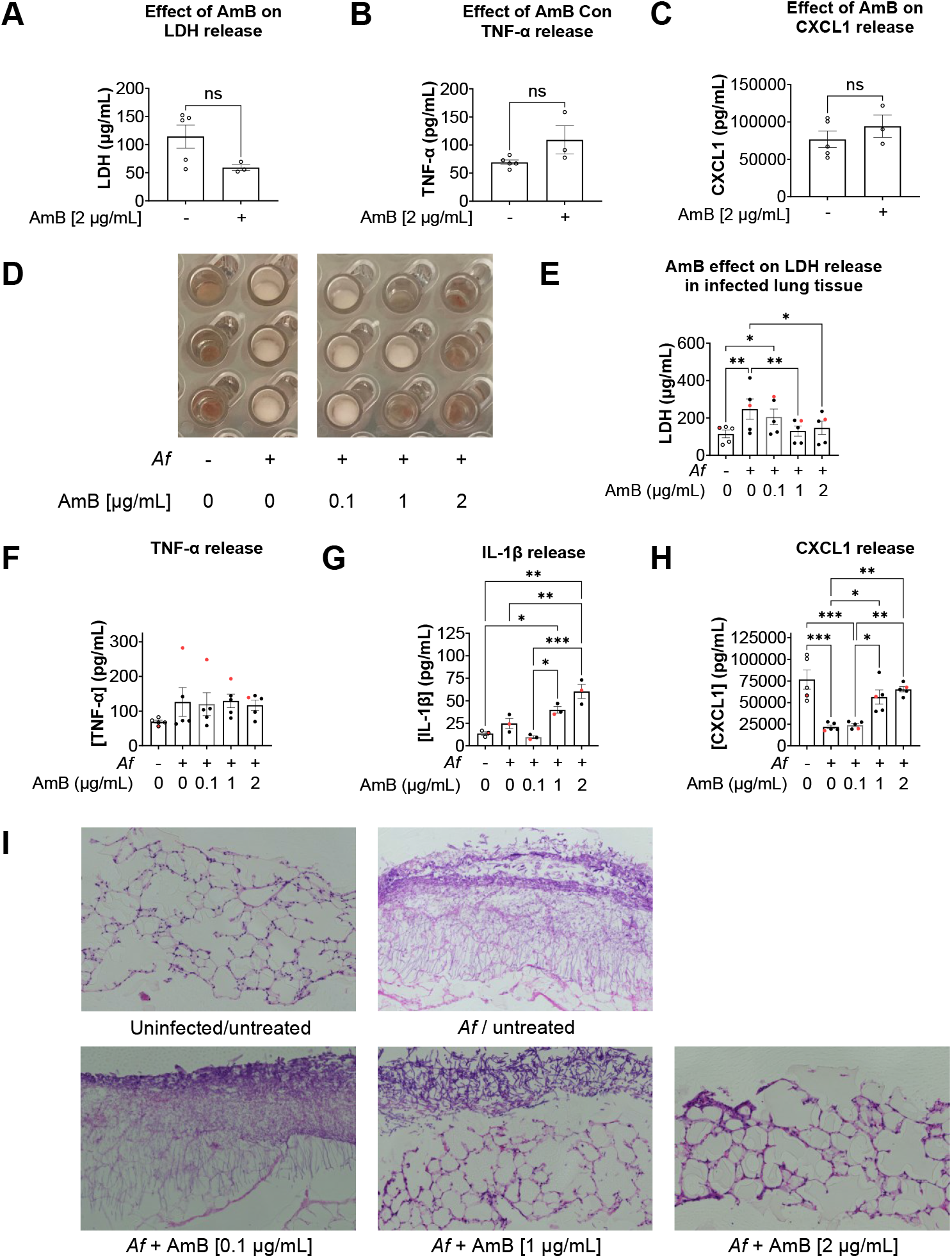
Amphotericin B (AmB) reduces tissue damage and modulates release of inflammatory cytokines in a dose-dependent manner. **A**. LDH released to supernatant from uninfected tissue discs treated for 48 hours with AmB [2 μg/mL]. **B**. Concentration of TNF-α in supernatant from uninfected tissue discs treated for 48 hours with AmB [2 μg/mL]. **C**. Concentration of CXCL1 in supernatant from uninfected tissue discs treated for 48 hours with AmB [2 μg/mL]. **D**. Photograph of lung discs on Transwell™ supports after 48 hours post infection showing effect of 3 different concentrations of Amphotericin B. **E**. LDH released to supernatant from infected lung tissue 48 hpi. **F**. TNF-α released 48 hpi. **G**. IL-1β released 48 hpi. **H**. CXCL1 released 48 hpi. **I**. Histology image (H&E) showing fungal growth and hyphae infiltration into the tissue with different concentrations off AmB. Data points in red represent values obtained from wells shown in panel D. One-way ANOVA on at least 3 biological replicates. p value * < 0.05, ** < 0.01, *** < 0.001, **** < 0.0001.

We investigated CXCL1 release further by measuring its concentration at 2 time-points, 24 and 48 hours. In uninfected lung discs, high levels of CXCL1 were detected at 24 hours and a second measurement at 48 hours showed that CXCL1 concentration doubled. Tissue infected with *Af* also showed high levels of CXCL1 at 24 hours; however, the second measurement at 48 hours showed that CXCL1 concentrations stagnated. The suppression of CXCL1 increase was reduced by AmB, which suggests *Af* has a role modulating tissue CXCL1 response (Fig. 5).

**Figure 5.**
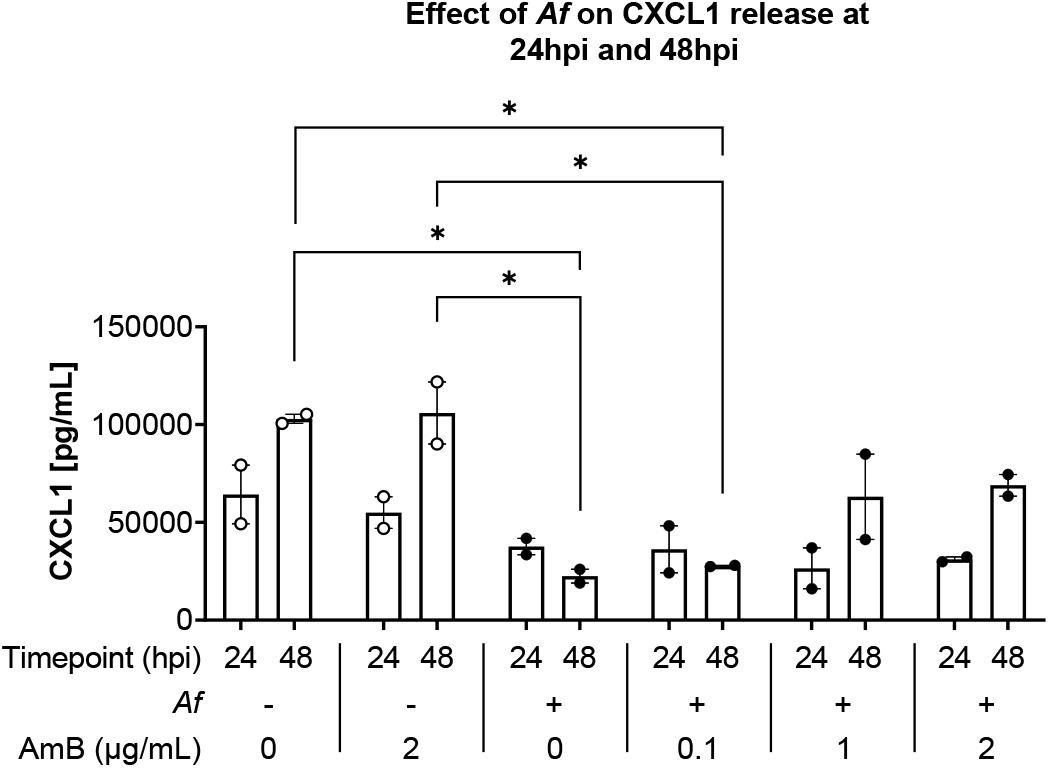
*Af* supresses increase of CXCL1 concentration in culture supernatant. Concentration of CXCL1 detected in supernatant at 24 hpi and 48 hpi. Technical triplicates per mouse were prepared from two animals. Statistical significance was tested with one-way ANOVA on biological replicates (n=2). p value * < 0.05, ** < 0.01, *** < 0.001, **** < 0.0001.

## DISCUSSION

Clinical data indicates that a subset of patients is susceptible to fungal disease. An association has been well established with neutropenia; however, the factors related to the diversity of clinical presentations, which include Allergic Bronchopulmonary Aspergillosis (ABPA), and Chronic Pulmonary Aspergillosis (CPA) associated with sarcoidosis or post-tuberculosis, are not thoroughly understood (1). We present an *ex-vivo* model focused on the characterization of pulmonary tissue responses to fungus and antifungal treatment. Each mouse can provide between 24 to 30 lung discs, allowing to test up to 9 conditions with three technical replicates. We propose as advantages of this model that it enables the study of different interventions in a single biological replicate, and the isolation of the pulmonary tissue, which helps to differentiate the immunological response triggered in the lungs without a systemic response contribution. Furthermore, these characteristics involve the fulfilment of the 3Rs’ principles of reduction and refinement (26).

Most experimental models, *ex-vivo* and *in-vivo*, for host-pathogen interactions are challenging to set-up with respect to control of important parameters associated with infection. *Af* adapts well to changes in the environment, which can lead to the modulation of its virulence. These adaptations are not exclusively dependent on nutrients, but also on conditions like pH and temperature (27,28). Transcriptome analysis has shown that *Af* changes with hypoxia by increasing the metabolism of polysaccharides, amino acids, metal ion transport, nitrogen and glycolysis (29) and these strategies seem to lead to enhanced invasion (30). An *in-vivo* model would require many experimental animals to control these parameters and *ex-vivo* models rely on submersion of tissue and fungi into culture medium, eliminating the air-liquid interface which has been shown to have immunological relevance. Immune responses to *Af* are mediated by alveolar macrophages and epithelial cells. Pathogen associated molecular patterns (PAMPs), like galactomannan and β-1,3-glucan, trigger initial inflammatory responses, the release of chemokines, cytokines, and promote cell migration to clear the fungi (31). Air-liquid interface has been shown to contribute significantly to epithelial cells function, enhancing surfactant, mucus production and modulating cytokines release (32).

To generate lung discs as homogeneous as possible, we applied the standard protocol of Placke et al. filling the lungs of C57BL/6J mice with 2% agarose. After the agarose solidified, the lungs were collected from the thoracic cavity, and immediately sliced. The functional residual capacity in a mouse is 0.25 mL, and we infused agarose to reach the total lung capacity of 1 mL (33), therefore pulmonary tissue represents only 25% of the final sample’s volume. For that reason, we included an overnight incubation step at 37°C before punching the lung discs. By removing the agarose, the lung slices become more compact, provide more tissue per volume of sample, and increases homogeneity between slices (Schematic 1).

Technically, an *ex-vivo* model based on PCLS implies variability in the proportions of the cellular components between lung discs. Minor histological differences can be expected based on the location of the lung from which the slice has been obtained. The closer the slice is to the hilum, the larger the calibre of the vessels and the airway. However, mouse lungs provide an advantage for this model, since the airway inside the lung parenchyma lacks cartilage (34), increasing uniformity between lung discs. However, perfect homogeneity between lung discs is impossible and variations could produce differences in the rate of nutrients consumption and medium acidification. Nonetheless, after 48 hours, one lung disc increases medium Abs at 415 nm on 59% (0.149) in comparison to medium only (0.094), keeping margin of error close to 5% (5.96% for CI=95%).

AmB showed an important immunomodulatory effect that agrees with previous research (35). Although, the effect on TNF-α did not reveal significant differences between conditions, a strong, dose-dependent increase in IL-1β was found. An interesting finding was the significant lower levels of CXCL1 from infected lung discs (Figure 4 F G H). Although the causes are not known, FBS has been shown to increase CXCL1 release in PCLS and some authors suggest to use serum-free media to avoid it (11); however, this approach reduces inflammatory cytokine responses (36).

AmB showed no effect on CXCL1 release on uninfected tissue (Fig 4 C) but had a dose-dependent effect to restore the release of CXCL1 by PCLS. This suggests that *Af* modulates directly CXCL1 responses. Further experiments are needed to elucidate the underlining mechanisms, with possible explanations including *Af* consuming the serum in the medium (37) reducing the stimulating factor for CXCL1 release. Alternatively, *Af* could reduce CXCL1 concentration through the activity of proteinases, that hydrolyse proteins.

Further analysis can be performed to study the variability of the cellular components between lung discs using flow cytometry. It has been shown that the proportion of some cell types, such as alveolar macrophages, can vary significantly between slices (11). Whether the variability reported of different ratios of alveolar macrophages influences TNF-α and IL-1β responses is not known and needs to be clarified. We decided to set a 48-hour time-point to ensure hyphal growth and confirm that this model enables infiltration into the tissue. Nonetheless, a limitation in this model is the difficulty to design a kinetic assessment of cytokine release, since large volumes of supernatant cannot be removed at multiple time-points from a single well.

Overall, the PCLS-ALI model reduces and refines the use of experimental animals for the investigation of pulmonary tissue immune responses, tissue damage, fungal invasiveness, and drug testing.

## ACKNOWLEDGEMENTS

We would like to thank Prof. Cecilia Johansson from the National Heart and Lung Institute, Imperial College London, for her expertise and assistance. Illustrations were created using BioRender.com.

## REFERENCES

1. Denning DW, Cadranel J, Beigelman-Aubry C, Ader F, Chakrabarti A, Blot S, et al. Chronic pulmonary aspergillosis: rationale and clinical guidelines for diagnosis and management. European Respiratory Journal. 2016 Jan 1;47(1):45–68.

2. Kosmidis C, Denning DW. The clinical spectrum of pulmonary aspergillosis. Thorax. 2015 Mar 1;70(3):270–7.

3. Vanden Bossche H, Warnock DW, Dupont B, Kerridge D, Sen Gupta S, Improvisi L, et al. Mechanisms and clinical impact of antifungal drug resistance. J Med Vet Mycol. 1994;32 Suppl 1:189–202.

4. Verweij PE, Ananda-Rajah M, Andes D, Arendrup MC, Brüggemann RJ, Chowdhary A, et al. International expert opinion on the management of infection caused by azole-resistant Aspergillus fumigatus. Drug Resist Updat. 2015 Aug;21–22:30–40.

5. Abdolrasouli A, Petrou MA, Park H, Rhodes JL, Rawson TM, Moore LSP, et al. Surveillance for Azole-Resistant Aspergillus fumigatus in a Centralized Diagnostic Mycology Service, London, United Kingdom, 1998-2017. Front Microbiol. 2018;9:2234.

6. Herbst S, Shah A, Carby M, Chusney G, Kikkeri N, Dorling A, et al. A new and clinically relevant murine model of solid-organ transplant aspergillosis. Dis Model Mech. 2013 May;6(3):643–51.

7. Buskirk AD, Templeton SP, Nayak AP, Hettick JM, Law BF, Green BJ, et al. Pulmonary immune responses to Aspergillus fumigatus in an immunocompetent mouse model of repeated exposures. Journal of Immunotoxicology. 2014 Apr 1;11(2):180–9.

8. Stolz DJ, Sands EM, Amarsaikhan N, Tsoggerel A, Templeton SP. Histological Quantification to Determine Lung Fungal Burden in Experimental Aspergillosis. J Vis Exp. 2018 Mar 9;(133):57155.

9. Dagenais TRT, Keller NP. Pathogenesis of Aspergillus fumigatus in Invasive Aspergillosis. Clin Microbiol Rev. 2009 Jul;22(3):447–65.

10. Franquet T, Müller NL, Giménez A, Guembe P, de la Torre J, Bagué S. Spectrum of Pulmonary Aspergillosis: Histologic, Clinical, and Radiologic Findings. RadioGraphics. 2001 Jul;21(4):825– 37.

11. Michalaki C, Dean C, Johansson C. The Use of Precision-Cut Lung Slices for Studying Innate Immunity to Viral Infections. Current Protocols. 2022;2(8):e505.

12. Viana F, O’Kane CM, Schroeder GN. Precision-cut lung slices: A powerful ex vivo model to investigate respiratory infectious diseases. Mol Microbiol. 2022 Mar;117(3):578–88.

13. Danov O, Lasswitz L, Obernolte H, Hesse C, Braun A, Wronski S, et al. Rupintrivir reduces RV-induced TH-2 cytokine IL-4 in precision-cut lung slices (PCLS) of HDM-sensitized mice ex vivo. Respir Res. 2019 Oct 22;20(1):228.

14. Zimniak M, Kirschner L, Hilpert H, Geiger N, Danov O, Oberwinkler H, et al. The serotonin reuptake inhibitor Fluoxetine inhibits SARS-CoV-2 in human lung tissue. Sci Rep. 2021 Mar 15;11(1):5890.

15. Choi KYG, Wu BC, Lee AHY, Baquir B, Hancock REW. Utilizing Organoid and Air-Liquid Interface Models as a Screening Method in the Development of New Host Defense Peptides. Front Cell Infect Microbiol. 2020;10:228.

16. Leroux MM, Hocquel R, Bourge K, Kokot B, Kokot H, Koklič T, et al. Aerosol-Cell Exposure System Applied to Semi-Adherent Cells for Aerosolization of Lung Surfactant and Nanoparticles Followed by High Quality RNA Extraction. Nanomaterials (Basel). 2022 Apr 15;12(8):1362.

17. He RW, Braakhuis HM, Vandebriel RJ, Staal YCM, Gremmer ER, Fokkens PHB, et al. Optimization of an air-liquid interface in vitro cell co-culture model to estimate the hazard of aerosol exposures. J Aerosol Sci. 2021 Mar;153:105703.

18. Amatngalim GD, Broekman W, Daniel NM, van der Vlugt LEPM, van Schadewijk A, Taube C, et al. Cigarette Smoke Modulates Repair and Innate Immunity following Injury to Airway Epithelial Cells. PLoS One. 2016 Nov 9;11(11):e0166255.

19. Perez-Cuesta U, Guruceaga X, Cendon-Sanchez S, Pelegri-Martinez E, Hernando FL, Ramirez-Garcia A, et al. Nitrogen, Iron, and Zinc Acquisition: Key Nutrients to Aspergillus fumigatus Virulence. J Fungi (Basel). 2021 Jun 28;7(7):518.

20. Dietl K, Renner K, Dettmer K, Timischl B, Eberhart K, Dorn C, et al. Lactic Acid and Acidification Inhibit TNF Secretion and Glycolysis of Human Monocytes. The Journal of Immunology. 2010 Feb 1;184(3):1200–9.

21. Jimdjio CK, Xue H, Bi Y, Nan M, Li L, Zhang R, et al. Effect of Ambient pH on Growth, Pathogenicity, and Patulin Production of Penicillium expansum. Toxins. 2021 Aug;13(8):550.

22. Pang KL, Chiang MWL, Guo SY, Shih CY, Dahms HU, Hwang JS, et al. Growth study under combined effects of temperature, pH and salinity and transcriptome analysis revealed adaptations of Aspergillus terreus NTOU4989 to the extreme conditions at Kueishan Island Hydrothermal Vent Field, Taiwan. PLOS ONE. 2020 May 26;15(5):e0233621.

23. Michl J, Park KC, Swietach P. Evidence-based guidelines for controlling pH in mammalian live-cell culture systems. Commun Biol. 2019 Apr 26;2(1):1–12.

24. Krycer JR, Lor M, Fitzsimmons RL, Hudson JE. A cell culture platform for quantifying metabolic substrate oxidation in bicarbonate-buffered medium. Journal of Biological Chemistry [Internet]. 2022 Feb 1 [cited 2022 Sep 5];298(2). Available from: https://www.jbc.org/article/S0021-9258(21)01357-0/abstract

25. Meletiadis J, Antachopoulos C, Stergiopoulou T, Pournaras S, Roilides E, Walsh TJ. Differential Fungicidal Activities of Amphotericin B and Voriconazole against Aspergillus Species Determined by Microbroth Methodology. Antimicrob Agents Chemother. 2007 Sep;51(9):3329– 37.

26. Hubrecht RC, Carter E. The 3Rs and Humane Experimental Technique: Implementing Change. Animals (Basel). 2019 Sep 30;9(10):754.

27. Hagiwara D, Sakamoto K, Abe K, Gomi K. Signaling pathways for stress responses and adaptation in Aspergillus species: stress biology in the post-genomic era. Bioscience, Biotechnology, and Biochemistry. 2016 Sep 1;80(9):1667–80.

28. Takahashi H, Kusuya Y, Hagiwara D, Takahashi-Nakaguchi A, Sakai K, Gonoi T. Global gene expression reveals stress-responsive genes in Aspergillus fumigatus mycelia. BMC Genomics. 2017 Dec 4;18(1):942.

29. Kroll K, Pähtz V, Hillmann F, Vaknin Y, Schmidt-Heck W, Roth M, et al. Identification of Hypoxia-Inducible Target Genes of Aspergillus fumigatus by Transcriptome Analysis Reveals Cellular Respiration as an Important Contributor to Hypoxic Survival. Eukaryot Cell. 2014 Sep;13(9):1241–53.

30. Hsu JL, Khan MA, Sobel RA, Jiang X, Clemons KV, Nguyen TT, et al. Aspergillus fumigatus invasion increases with progressive airway ischemia. PLoS One. 2013;8(10):e77136.

31. Arias M, Santiago L, Vidal-García M, Redrado S, Lanuza P, Comas L, et al. Preparations for Invasion: Modulation of Host Lung Immunity During Pulmonary Aspergillosis by Gliotoxin and Other Fungal Secondary Metabolites. Frontiers in Immunology [Internet]. 2018 [cited 2022 Dec 19];9. Available from: https://www.frontiersin.org/articles/10.3389/fimmu.2018.02549

32. Silva S, Bicker J, Falcão A, Fortuna A. Air-liquid interface (ALI) impact on different respiratory cell cultures. European Journal of Pharmaceutics and Biopharmaceutics. 2023 Mar 1;184:62–82.

33. Lai YL, Chou HC. Respiratory mechanics and maximal expiratory flow in the anesthetized mouse. Journal of Applied Physiology. 2000 Mar;88(3):939–43.

34. Meyerholz DK, Sieren JC, Beck AP, Flaherty HA. Approaches to Evaluate Lung Inflammation in Translational Research. Vet Pathol. 2018 Jan 1;55(1):42–52.

35. Mesa-Arango AC, Scorzoni L, Zaragoza O. It only takes one to do many jobs: Amphotericin B as antifungal and immunomodulatory drug. Frontiers in Microbiology [Internet]. 2012 [cited 2022 Nov 30];3. Available from: https://www.frontiersin.org/articles/10.3389/fmicb.2012.00286

36. Lam M, Lamanna E, Organ L, Donovan C, Bourke JE. Perspectives on precision cut lung slices—powerful tools for investigation of mechanisms and therapeutic targets in lung diseases. Frontiers in Pharmacology [Internet]. 2023 [cited 2024 Jan 24];14. Available from: https://www.frontiersin.org/articles/10.3389/fphar.2023.1162889

37. Gifford AHT, Klippenstein JR, Moore MM. Serum stimulates growth of and proteinase secretion by Aspergillus fumigatus. Infect Immun. 2002 Jan;70(1):19–26.

